# MAGE: Strain Level Profiling of Metagenome Samples

**DOI:** 10.1101/2022.11.24.517382

**Authors:** Vidushi Walia, V.G Saipradeep, Rajgopal Srinivasan, Naveen Sivadasan

## Abstract

Metagenomic profiling from sequencing data aims to disentangle a microbial sample at lower ranks of taxonomy, such as species and strains. Deep taxonomic profiling involving accurate estimation of strain level abundances aids in precise quantification of the microbial composition, which plays a crucial role in various downstream analyses. Existing tools primarily focus on strain/subspecies identification and limit abundance estimation to the species level. Abundance quantification of the identified strains is challenging and remains largely unaddressed by the existing approaches. We propose a novel algorithm MAGE (Microbial Abundance GaugE), for accurately identifying constituent strains and quantifying strain level relative abundances. For accurate profiling, MAGE uses read mapping information and performs a novel local search-based profiling guided by a constrained optimization based on maximum likelihood estimation. Unlike the existing approaches that often rely on strain-specific markers and homology information for deep profiling, MAGE works solely with read mapping information, which is the set of target strains from the reference collection for each mapped read. As part of MAGE, we provide an alignment-free and kmer-based read mapper that uses a compact and comprehensive index constructed using FM-index and R-index. We use a variety of evaluation metrics for validating abundances estimation quality. We performed several experiments using a variety of datasets, and MAGE exhibited superior performance compared to the existing tools on a wide range of performance metrics.

## 1 Introduction

The dynamics of a microbial ecosystem is governed by the diversity and abundance of microorganisms present in the microbial environment. Metagenomics research empowers the detailed characterization of microbial population in the microbial environment or host-associated microbial communities. The detailed analysis of microbial communities in metagenomic samples is tremendously useful in clinical study, disease study, study of environmental habitats, characterization of an industrial product, understanding the host-pathogen interactions and many more. Metagenomic sequencing [27] is a powerful way of examining the diversity and complexity of microbial communities. Advancements in Next Generation Sequencing (NGS) technologies have greatly facilitated culture independent analysis of the microbiome, which has increased the scope of metagenomic studies [12, 2].

An essential prerequisite for any metagenomic analysis is to disentangle the microbial sample at lower ranks of taxonomy such as species/strain with precise measurements of their abundances. Within the same species individual strains can elicit different functions and metabolic responses [3, 1], thus governing the dynamics of the microbial environment. This establishes strains as the functional unit of microbial taxonomy. This necessitates identification and quantification of metagenomic samples down to the level of strains.

Characterizing or profiling of metagenomic samples at the level of strains becomes even more complicated due to the high similarity of the genomes sequences present in the samples. Selective amplification of highly conserved regions is widely used for microbial profiling. These conserved regions include housekeeping genes such as 16s and 18s rRNA, ITS. 16srRNA has been extensively used by the microbial community. However, it is not only incapable of capturing the viruses, plasmids and eukaryotes present in the microbial community, but also has a low discriminating power for the classification at species and strain level [19]. Other conserved gene regions or the repetitive regions which have high evolutionary rate offer improved classification at species and strain level but suffer from underdeveloped reference collections [19] and low accuracy in estimating the abundances at strain-level. Whole genome sequencing approaches on the other hand are promising and address most of the drawbacks of amplicon based approaches but suffer from the problems of high cost and shorter read length which results in ambiguous mapping. There are several strategies to accomplish metagenomic profiling which includes alignment/mapping of reads to complete reference sequences or to a well curated database of unique taxonomic markers, k-mer composition based approaches and assembly based approaches. The approaches based on unique taxonomic markers work well in discriminating strains in microbial samples but lacks accuracy in estimating their abundance. These approaches do not allow for easy updates in the reference database as they rely on complex and compute intensive methods to build reference collection of signature markers for every level of taxonomy.

There is a plethora of microbial abundance estimation tools [19] [27] that address the problem of species level profiling and strain level identification. However, problem of strain level abundance estimation remains challenging and largely unaddressed. Tools like MetaPhlAn_strainer [29, 29, 4] provides strain tracking feature along with abundance estimation at strain level. StrainPhlAn2 [4] solves a different problem of SNP profiling of strains present in a metagenomic sample. PanPhlAn [26] and StrainPhlAn2 both work at the strain-level resolution. StrainPhlAn2 provides the SNP profile of the detected strains and PanPhlAn provides the gene specific information for the strains present in the sample. Other approaches like GOTTCHA [10] (a gene-independent, signature-based approach) and Centrifuge [14] (a high speed metagenomic classifier), provide more detailed taxonomic profiles which includes abundance estimates at the strain-level.

Kraken2 [30, 31] solves the problem of sub-species/strain level identification and relies on Bracken [18] for efficient estimation of species-level microbial abundance. Kraken2 also reports the number of reads assigned to a taxa in a metagenomic sample, this information can be further utilized to estimate the abundance at various levels of the taxonomy. Other approaches like StrainGE [8], Strain-FLAIR [7] provides strain level resolution. However, these approaches work with a very limited reference collections and thus are not suitable as general purpose profiling tools. For example StrainGE [8] provides strain-level profiling but limit the analyses to within species strains in bacterial population and focuses on strain-aware variant calling.

We develop a novel algorithm MAGE (Microbial Abundance GaugE), for accurately identifying constituent strains and quantifying strain level relative abundances. For accurate and efficient profiling, MAGE uses read mapping information and performs a novel local search-based profiling guided by a constrained optimization based on maximum likelihood estimation. Unlike the existing approaches that often rely on taxonomic markers and homology information for deep profiling, MAGE works solely with read mapping information, which is the set of target strains from the reference collection for each mapped read. As part of MAGE, we provide an alignment-free and kmer-based read mapper (MAGE mapper) that uses a compact and comprehensive index constructed using FM-index [9] and R-index [11]. In MAGE mapper, the reads are mapped to strains solely based on the *k*-mer composition of the reads without requiring any gapped alignment. We use a variety of evaluation metrics for validating abundances estimation quality. We benchmark MAGE against state-of-the-art methods using a variety of low complexity and high complexity datasets. MAGE showed superior performance compared to the state-of-the-art on a wide range of performance measures.

## 2 Methods

In this section, we discuss details of MAGE abundance estimation and MAGE mapper. First we discuss how MAGE estimates strain level abundances given read mapping information. In the later sections, we present an alignment free and *k*-mer based mapper to generate read mapping information.

### 2.1 Abundance Estimation and MLE

Let *S* = {*s*_1_, …, *s*_*N*_} denote a reference collection of *N* strains (sequences) spanning the various members of the microbial community such as bacteria, virus, fungi etc. We will use the terms sequence and strain interchangeably. For strain level abundance estimation, MAGE uses read mapping information, which is the set of target strains from the reference collection *S* for each mapped read. Let ℛ denote the set of mapped reads. For a read *r* ∈ ℛ, let *Q*(*r*) ⊆ *S* denote the subset of strains from *S* to which read *r* maps. The read mapping information is simply {*Q*(*r*)}. In the following we first discuss how MAGE estimates strain level abundances given {*Q*(*r*)}.

Before discussing the details of the approach used by MAGE, we first discuss the standard MLE (Maximum Likelihood Estimation) formulation similar to [5, 25] for abundance estimation. Each read *r* in ℛ is assumed to be sampled independently and randomly from its source strain. The source strain is chosen with probability *a*_*i*_, which is the relative abundance of *s*_*i*_ in the sample. We make the simplifying assumption of uniform coverage, which implies that a read is generated from a location sampled uniformly at random from its source strain. Let 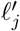 denote the effective length of strain *s*_*j*_ [5, 25]. Under the uniform coverage assumption, we approximate 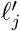 as 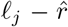 where 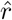 is the average read length. More accurate estimates for 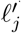 can also be used [25]. For a given sample, the relative abundances of all the strains is denoted by the vector **a** where the *i*th component denotes the relative abundance of the *i*th reference strain. Clearly, ∑*a*_*i*_ = 1. The likelihood for ℛ is given by

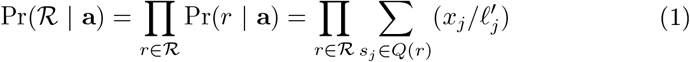

where *x*_*j*_ and *a*_*j*_ are related as 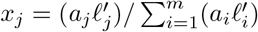. Here, *x*_*j*_ is simply the length adjusted relative abundance of strain *s*_*j*_. Clearly, ∑*x*_*i*_ = 1. For several reads, their corresponding sequence sets *Q*(*r*) will be identical. In other words, the mapped sequence sets of the reads define a partition of ℛ = 𝒞_1_ ∪ … ∪ 𝒞_*R*_ where *C*_*i*_ denotes the subset of reads that are mapped to the same set of sequences denoted as *Q*_*i*_. Therefore, the log likelihood with respect to the strain collection *S* is given by

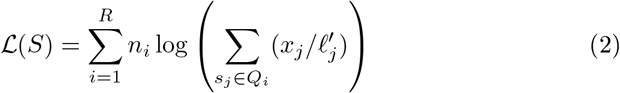

where *n*_*i*_ is the cardinality of 𝒞_*i*_. We solve for *x*_*j*_s that maximizes ℒ. From *x*_*j*_, we solve for *a*_*j*_ using their relation. To reduce estimation noise, estimated abundance values below a small predefined threshold can be made zero.

The standard MLE approach usually works well for species level abundance estimation. However, MLE suffers from the problem of distributing abundance values across several related strains that are present in the reference leading to reduced specificity and strain level profiling quality. This is because, inter-strain sequence similarity within a species is much higher than inter-species similarity.

### 2.2 Constrained Optimization and Strain Level Coverage

To perform accurate strain level profiling, MAGE performs a local search based profiling guided by the following constrained optimization problem based on the MLE formulation discussed earlier. EM based solution of the above MLE formulation also yields an estimate of the latent matrix *M*_*R×N*_, where *M* (*i, j*) is the number of reads from the read partition 𝒞_*i*_ that have originated from strain *s*_*j*_. We recall that all reads in 𝒞_*i*_ map to the same subset of strains *Q*_*i*_. If strain *s*_*j*_ does not belong to *Q*_*i*_ then clearly *M* (*i, j*) = 0. From an optimal MLE solution, estimate for *M* (*i, j*), where strain *s*_*j*_ belongs to *Q*_*i*_, is obtained as

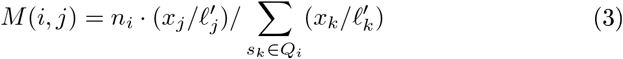

where we recall that *n*_*i*_ is the total number of reads in the partition 𝒞_*i*_ and *Q*_*j*_ is the set of sequences to which each read in 𝒞_*i*_ maps to. Using matrix *M*, estimate for the total number of reads originating from strain *s*_*j*_ denoted by *Y*_*j*_ is given by

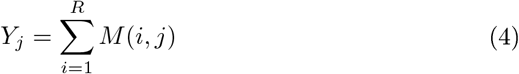

Also, *Y*_*j*_ and *x*_*j*_ are related as

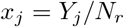

where *N*_*r*_ = ∑ *n*_*i*_ = |ℛ| is the total number of reads in ℛ.

Using the estimate of *Y*_*j*_, estimated coverage of strain *s*_*i*_, denoted by *cov*(*j*), is given by

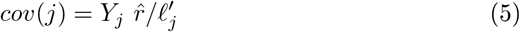

where we recall that 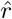 is the average read length and 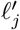 is the effective length of strain *s*_*j*_. From the above set of equations, we obtain that optimal estimates for *x*_*j*_ and *cov*(*j*) are related as

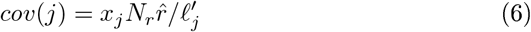

Coverage values *cov*(*j*) for different strains can vary as it depends on both read coverage as well as the abundance of the strain in the sample. Using eq (6) and noting that *N*_*r*_ and 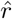 are constants, it is straightforward to verify that at an optimal MLE solution for (2), the objective value ℒ° is given by

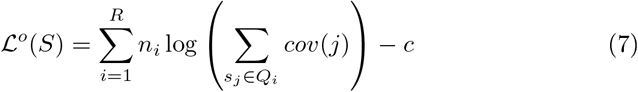

where the 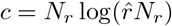 is a constant. From the above expression, we see that the final optimum value depends on the cumulative coverages of the strain sets *Q*_1_, …, *Q*_*R*_ where the cumulative coverage of set *Q*_*i*_ is the sum total of the strain coverages of strains in *Q*_*i*_.

Suppose we wish to perform a constrained MLE where *S* is restricted to a subset say *S*′ ⊂ *S*. When *S*′ is specified, the original read mapping, namely, *Q*_1_, …, *Q*_*R*_, undergoes a projection to obtain the modified mapping information 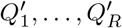 where 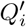 is simply 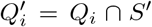. That is, only the strains in *S*′ are retained in 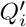. In the rest of the paper, when a strain subset *S*′ is specified, we assume without explicitly stating that the associated mapping information 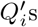 are the projections of *Q*_*i*_s obtained with respect to *S*. It is easy to see that restricting the strain set to *S*′ amounts to constraining remaining strains to have zero abundance. MLE with respect to *S*′ has its optimum value again given by ℒ°(*S*′).

In the next section, we discuss how MAGE performs strain set refinement using a local search, guided by strain coverages *cov*(*j*) and ℒ^*o*^(*S*′), in order to improve the specificity and profiling accuracy. We note that under strain set constraints, for a strain *s*′ ∈ *S* − *S*′, the reads that are estimated to be from strain *s*′ in the unconstrained MLE are redistributed to those strains in *S*′ that occur along with *s*′ in any of the original strain subset *Q*_*i*_. This redistribution leads to an increase in the estimated coverages of the strains in *S*′. From ℒ^*o*^(*S*′), we observe that the final optimum objective value depends not only on the individual coverages *cov*(*j*) of the strains in *S*′ but more importantly the cumulative coverages with respect to the associated 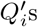. Hence, strain set restriction should be guided in a manner that increases the cumulative coverages of associated 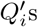.

One additional aspect to be considered for strain set refinement is that when the strain set *S* is restricted to *S*′, some of the reads in ℛ can become unmapped because none of the mapped strains for this read are present in *S*′. This is a low probability event if *S*′ contains all the strains present in the sample. Let *p*_*ϵ*_ denote the probability of observing such an unmapped read. From the independence assumption, the probability of observing *t* such reads is 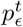. As a consequence, in the likelihood expression given by eq (1), each unmapped read contributes a multiplicative term *p*_*ϵ*_. Lastly, an additional sparsity inducing regularizer is introduced in the objective function to favour sparse solutions. Consequently, the optimum objective value ℒ° in eq (7) is extended as

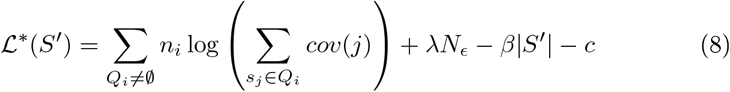

where *S*′ is the restricted set, *Q*_*i*_s are the projected strain sets with respect to *S*′, *λ* = log(*p*_*ϵ*_), *β* is the sparsity parameter, and *N*_*ϵ*_ is the number of unmapped reads with respect to *S*′. In the next section, we discuss how MAGE performs iterative strain set refinement using a local search guided by the above constrained objective function.

### 2.3 Local Search and Strain Set Refinement

Suppose we know the true set of strains *S*′. Then the abundance estimation can be done by simply performing the constrained MLE (ℒ^*o*^(*S*′)) as discussed in the previous section. However, in practice we do not have such knowledge. Estimating ℒ*(*S*′) for all possible subsets is also prohibitively expensive as there are exponentially many subsets. Additionally, if we have prior information on lower bounds for strain level coverages (*cov*(*j*)), we could further prune the search space to those *S*′ for which the estimated coverages satisfy the additional constraints. However, this information is also hard to obtain, because, as discussed earlier, coverages *cov*(*j*) depends on the strain abundances and coverage induced by the mapped reads, which can deviate due to strain mutations. Further, the regularizers used in the above objective do not exploit any specific priors on the cardinality of the distinct strains in the given sample or its composition. Because of this, a set corresponding to an optimal solution for the above objective can still contain several false positives. MAGE therefore uses the above objective ℒ* as a surrogate for the true unknown objective. MAGE performs a local search based iterative strain set refinement guided by the surrogate objective ℒ* in order to arrive at its final strain set *S*′ which is close to the (unknown) true set. In the following, we discuss the the iterative strain set refinement performed by MAGE.

Starting with the initial MLE solution on the on the full strain set *S*, MAGE iteratively removes strains from among the candidates that have low estimated coverages. The details are discussed later. Removal of such strains can induce only minimal changes to ℒ^*o*^(*S*′) in each iteration as there are only few reads that gets redistributed in each iteration. Therefore, performing refinement in this manner helps the local search and refinement not to deviate from the desired trajectory in the solution space that leads to the true final *S*\*.

To start with, all strains whose coverage is below a low threshold are removed at once before performing the iterative refinement. In each iteration of strain set refinement, greedily removing the strain with the lowest estimated coverage can lead to false negatives. This is because, some of the true strains can have low estimated coverages in the current solution *S*′ due to the presence of several similar false positive strains in *S*′. MAGE therefore considers the bottom *b* strains 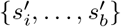 having the lowest estimated coverages *cov*(*j*), for some fixed *b*, and computes the modified 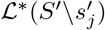 values for each of these *b* candidates. Among these *b* candidates, we finally remove, from the current solution, the strain *s*′ whose corresponding ℒ*(*S*′ \*s*′) is the largest. The refined set after this iteration is given by *S*′ \*s*′. The coverage values of the strains in the refined set are updated using the outcome of MLE performed (using eq(5)) for estimating ℒ*(*S*′ \*s*′).

To reduce the overall number of MLE computations, MAGE further approximates 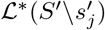 for a low coverage strain 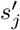 as follows. The current sets *Q*_*i*_ are first projected with respect to 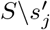 as discussed earlier. Using the projected 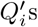 and using eq (3), *M* (*i, j*) values are directly recomputed. Using these updated *M* (*i, j*) values, modified coverage values *cov* (*j*) for the strains in 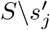 are computed using equations (4) and (5). These modified coverage values and the modified count of unmapped reads 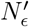 (as a consequence of removing 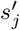) are used in eq (7) to approximate ℒ*(*S*\*s*_*j*_). The above approximation is performed for every *t* consecutive iterations, for a fixed *t*, followed by MLE and computation of ℒ*(*S*′) without approximation in the subsequent iteration.

As discussed earlier, after each iteration, the coverages *cov*(*j*) of the remaining strains in the solution set *S*′ gradually increase. However, excessive refinement will lead to significant decline in the ℒ*(*S*′) value because of removing true positives and thereby leading to several unmapped reads. MAGE identifies the final subset *S*′ in the following manner. MAGE first performs the above strain filtering operation till no more strains are left. This produces a sequence of strains *s*_1_, …, *s*_*m*_ in the order of their removal, where the strain *s*_*m*_ in the end of this sequence was removed in the last filtering iteration *m*. We expect that the true positive strains are present in abundance towards the end of this sequence. In other words, the rank of a strain in the above sequence in a sense indicates the likelihood of the strain to be present in the sample. Let the change of the objective function value at each of these *m* iterations be denoted by the corresponding sequence *δ*_1_, …, *δ*_*m*_. Using this sequence of *δ*s, MAGE attempts to identify an iteration *k* ≤ *m* such that the final strain set *S* given by the sequence *s*_*k*_, …, *s*_*m*_ has reduced number of false positives and false negatives. For computing *k*, MAGE performs change point detection on the sequence of *δ*s where the change point indicates the iteration after which significant number of true positives are filtered. The change point is detected using a simple heuristic: scan in the order *δ*_*m*_, …, *δ*_1_ and consider the corresponding moving averages 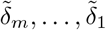 where 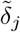 is the average of *δ*_*j*_, *δ*_*j*+1_ and *δ*_*j*+2_. Stop at *k* such that 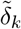 is more than a threshold *Δ* and 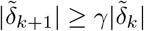 for a threshold *γ*. For the final strain set *S*′, the associated abundance values are computed using MLE as discussed earlier.

### 2.4 Read Mapping

As part of MAGE, we provide an alignment-free and kmer-based read mapper to compute the read mapping information {*Q*(*r*)} which forms the input for the abundance estimation discussed above. The read mapper works solely based on *k*-mer lookup on the reference collection. That is, for each *k*-mer in the read *r*, the target set of strains in the reference where the *k*-mer is present is identified. After querying all *k*-mers in the read, we finally obtain a candidate subset of strains *C* from the reference that received one or more *k*-mer hits. The final set of strains *Q*(*r*) that qualify as target strains for the mapped read *r* is identified by performing a random selection from the candidate strain set *C*. The random selection is based on the cumulative *k*-mer hits that a strain in *C* has received. For this, MAGE maintains a probability distribution that encodes Pr(cumulative *k*-mer hits = *t* for a read on its source strain). This distribution is estimated apriori using read simulations. Using this probability distribution, each candidate strain in *C* is included in *Q*(*r*) based on the probability associated with the cumulative *k*-mer hits for the strain.

## Reference Index

To support *k*-mer lookup required by the MAGE mapper, we also constructed an index of the reference collection. We remark that any index of the reference that supports *k*-mer search would suffice for the MAGE read mapper. However, as part of MAGE, we constructed a simple index by full text indexing of the reference using FM-index [9] and R-index [11]. R-index provides superior compression when the constituent strains share high similarity. However, for divergent set of strains, R-index exhibits poor compression compared to FM-index. We therefore used R-index to index the sub-collection of species that have several strains with high inter-strain similarity. For the remaining species, we used FM-index. In order to link the sub-collection indexes, we created an additional meta-index that directs the *k*-mer queries to the appropriate sub-collection indexes. The meta-index is a simple lookup that maps a *k*-mer to a binary vector where a bit is ON iff the *k*-mer is present in the associated sub-collection. Clearly, the above meta-index allows the reference to be chunked if required into several sub-collections. However, we remark that the meta-index is optional. For instance, no additional meta-index is required when the whole reference indexed as a single chunk. We used the Refseq strain collection [23] as the reference in the current implementation and the associated index that we created used meta-index. For a given read, all the read *k*-mers are first searched only in the meta-index. Only those index chunks that received sufficiently many *k*-mer hits, parameterized by a threshold *h*, are subsequently searched for the candidate strains.

## 3 Results

We compare MAGE against state-of-the-art metagenome strain level profiling tools Centrifuge [14], GOTTCHA [10], Kraken2 [31] and MetaPhlAn_strainer [29].

For MetaPhlAn_strainer, we used the tool accessed from [21], the strain tracking feature of MetaPhlAn_strainer enables profiling the samples down to the resolution of strains. MetaPhlAn_strainer [29] database has extended set of markers which enables strain identification and strain tracking of metagenomic samples. StrainPhlAn2 and PanPhlAn do provide strain level relative abundances. We therefore chose MetaPhlAn_strainer over the other tools offered by bioBakery [20] in our benchmark. Centrifuge recommends uncompressed index for taxonomic profiling at the strain level [14]. Hence, we considered only those indexes that are constructed from uncompressed genomic sequences. For benchmarking MAGE’s results with Centrifuge, we have considered two different indexes provided by Centrifuge, which we denote as Centrifuge1 and Centrifuge2 in the results. Centrifuge1 uses “p+h+v” index for read mapping. It includes reference genomes from Bacteria, Archae, Viruses and Human (2016 release). Centrifuge2 uses the massive “nt-database” (compressed index size 64GB) index built using NCBI nucleotide non-redundant sequences (2018 release). These indexes are available for download from [6]. GOTTCHA also provides multiple index for metagenomic profiling. In the results GOTTCHA 1 and GOTTCHA 2 denote the results from the two indexes that we have considered in our benchmarking. GOTTCHA 1 uses bacterial index “GOTTCHA_BACTERIA_c4937_k24_u30_xHUMAN3x.strain” for profiling metagenome samples while GOTTCHA 2 uses the combined index “GOTTCHA_COMBINED_xHUMAN3x.strain” for profiling. Both GOTTCHA indexes selected for benchmarking offer strain-level resolution.

Kraken2 also offers two mini databases, namely MiniKraken version1 and MiniKraken version2, that are constructed by downsampling the minimizers in the standard Kraken2 database. The sole difference between the two Minikraken databases is that MiniKraken version2 also includes kmers from GRCh38 human genome. The results for these two different databases is are denoted as Kraken V1 and Kraken V2 respectively. Since Kraken2 does not report the relative abundances directly, we estimate the strain level abundances as relative fraction of the read counts (reported by Kraken2) after normalizing by the corresponding genome length. Since all ground truth strains belong to Refseq, we considered only the Refseq strains reported by Kraken2.

The strain level outputs of MetaPhlAn_strainer, GOTTCHA 1&2, Kraken2 V1&V2 and Centrifuge 1&2 were mapped to the NCBI taxonomy to compare them with the ground truth. For all experiments, the MAGE parameters were fixed as 21 for *k*-mer length in level 2 index, 16 for *k*-mer length in level 1 index and 3 for parameter *h* used in read mapping. In the abundance estimation phase, the parameter values used were *b* = 5, *t* = 5, *β* = 2000, *γ* = 2, *Δ* = 10^6^.

We additionally included the standard MLE solution (described in the Section 2.1) in our benchmarking. For the standard MLE, the pruning threshold for relative abundance was set to 10^−4^.

### 3.1 Reference Collections

MAGE index was constructed using reference genome sequences from NCBI’s Refseq (complete genomes) [23], that were available at the time of download (2018). In total, MAGE reference database consisted of ~24k strains spanning ~13k species. The species were from 3,559 genus belonging to Virus, Bacteria, Archae, Fungi and Protozoa. The final MAGE index was ~38 GB in size. The level 2 MAGE index consisted of 32 chunks of which 22 were indexed using R-index and the remaining 10 were indexed using FM-index.

The Refseq strains were part of the reference collection used by each of the tool used in our benchmarking. MetaPhlAn_strainer uses reference genomes from Integrated Microbial Genomes (IMG) system to build its database of clade-specific marker genes. The IMG database contains sequence data from multiple sources including Refseq.

The “nt-database” index of Centrifuge [14] used by Centrifuge2 contains prokaryotic sequences from NCBI BLAST’s nucleotide database which contains all spliced non-redundant coding sequences from multiple database including Refseq. The “p+h+v” Centrifuge index used by Centrifuge1 consists of prokaryotic and human genome sequences downloaded from NCBI and contains sequences from Bacteria, Archae, Viruses and Human.

GOTTCHA provides a database of unique signatures for prokaryotic and viral genomes. These unique genome segments are scattered across multiple levels of taxonomy. Minikraken indexes version1 and version2 used by Kraken2 V1 and V2 respectively also incorporate reference genomes from Refseq.

### 3.2 Datasets

We experimented with multiple datasets, each containing strains from across the various kingdoms of prokaryotes. We used ART [13] simulator to generate Illumina paired-end reads of 150bp length and with 20X coverage, to generate the various synthetic read datasets.

We created three simulated read datasets, namely, HC1, HC2 and LC. Both HC1 and HC2 datasets are high complexity metagenomic samples, where multiple strains from same species or species with same genus are present in a sample, such that the constituent strains bear high similarity.

HC1 dataset consisted of 34 strains from 10 species spanning 8 genus. Out of the 34 strains, 20 strains were from Escherichia Coli and the remaining 14 strains were spread across 9 other species. Of the 10 species in HC1, 9 species were from bacteria and the remaining 1 was from a viral species (“Murine leukemia”). In HC1 sample, most of the constituent strains had similar relative abundances and a very few strains had higher abundances.

The HC2 dataset consisted of 60 strains from 25 species spanning 25 genus. HC2 dataset had 5 strains from Salmonella enterica, 4 strains from “Bordetella pertussis”. The remaining 23 species had 2 to 3 strains each. Out of the 25 species, 24 species are from Bacteria and 1 was a viral species (“Bixzunavirus Bxz1”). In HC2 sample, the constituent strains had varying relative abundances.

LC is a low complexity dataset where most constituent strains do not bear very high similarity with one another. The LC dataset consisted of 50 strains from 47 species in total spanning 38 genus. Each of the 47 species contributed 1 to 2 strain to the LC sample. In LC sample, close to 20% of the strains had higher relative abundances and the remaining 80% of the strains had similar and comparatively lower relative abundances.

To mimic the real world metagenomic samples, we also constructed simulated read datasets where the reference strains were included in the sample after introducing mutations. This allows us to evaluate the ability of the benchmarked tools to correctly identify and quantify the target strains in the presence of mutations. In this study, we used three datasets, namely HC1-Mutated, HC2-Mutated and LC-Mutated, after introducing mutations to the strains in HC1, HC2 and LC respectively. We used wgsim [17, 16] pipeline to create the read datasets with 20X coverage. Wgsim introduces SNPs and insertion/deletion (INDEL) mutations to the input genomes and simulates reads from the mutated genomes. We used default parameters to generate mutated read dataset with 0.1% mutation, of which 15% of the mutations were INDELS and the remaining 85% mutations were SNPs. For HC1, HC2 and LC, mutations were introduced in all the strains present in the sample.

### 3.3 Performance Metrics

We used a wide variety of metrics to benchmark the various tools used in our experiments. We used four well-known divergence measures for probability distributions to compare relative abundance distributions, the Jensen-Shannon divergence(JSD), Total variation distance(TVD), Hellinger distance(HD) and cumulative mass (CM). The Jensen-Shannon divergence (JSD) between two distributions *P* and *Q*, which is a smoothed and symmetric version of the KL-divergence [28], is given by *JSD*(*P* ∥ *Q*) = *KL*(*P* ∥ *M*) + *KL*(*Q*∥ *M*) where 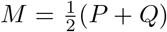. Square root of JS divergence is known to be a metric. Total variation distance (TVD) *δ*(*P, Q*) between distributions *P* and *Q* is a statistical distance metric and is given by 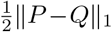. Hellinger distance [22] *H*(*P,Q*) between distributions *P* and *Q* is given by 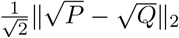, where 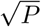 denotes the component-wise square root of *P*. Hellingner distance is known to be a metric and is also related to Bhattacharya coefficient. JSD, TVD and HD are known to be special cases of *f*-divergences between distributions. The cumulative mass (CM) of the true positives for an estimated distribution *P* with respect to a true distribution *Q* is given by the sum 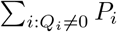. Additionally, we also compare the ability of the tools to correctly identify strains using the standard precision, recall and F1-score metrics. For JSD, TVD and HD, lower values indicate better performance. For CM, precision, recall and F1-scores, higher values indicate better performance.

### 3.4 Results on HC1, HC2 and LC Datasets

Table 1, Table 2 and Table 3 shows the comparison of the benchmarked tools on the high complexity datasets HC1 and HC2 and the low complexity data LC. For all experiments, the best score under each metric is highlighted in bold.

**Table 1.** Strain level profiling performances on HC-1 data

|  | MetaPhlAn<br>strainer | Centrifuge<br>1 | Centrifuge<br>2 | GOTTCHA<br>1 | GOTTCHA<br>2 | Kraken2<br>V1 | Kraken2<br>V2 | MLE | MAGE |
| --- | --- | --- | --- | --- | --- | --- | --- | --- | --- |
| **JSD** | 0.8197 | 0.7468 | 0.8305 | 0.6960 | 0.7496 | 0.6928 | 0.6950 | 0.2829 | **0.2287** |
| **TVD** | 0.9714 | 0.8515 | 0.9988 | 0.7714 | 0.8523 | 0.8138 | 0.8159 | 0.1721 | **0.0974** |
| **HD** | 0.9845 | 0.8892 | 0.9923 | 0.8312 | 0.8981 | 0.7902 | 0.7931 | 0.3315 | **0.2739** |
| **CM** | 0.0331 | 0.1888 | 0.0012 | 0.4281 | 0.1488 | **0.9608** | 0.9495 | 0.8381 | 0.9377 |
| **Prec.** | 0.1111 | 0.0323 | 0.0233 | 0.0833 | 0.0606 | 0.0102 | 0.0100 | 0.2063 | **0.9118** |
| **Recall** | 0.0294 | 0.3235 | 0.2941 | 0.2059 | 0.2353 | 0.2353 | 0.2353 | 0.9706 | **0.9118** |
| **F1** | 0.0465 | 0.0587 | 0.0431 | 0.1186 | 0.0964 | 0.0195 | 0.0192 | 0.3402 | **0.9118** |

**Table 2.** Strain level profiling performances on HC-2 data

|  | MetaPhlAn<br>strainer | Centrifuge<br>1 | Centrifuge<br>2 | GOTTCHA<br>1 | GOTTCHA<br>2 | Kraken2<br>V1 | Kraken2<br>V2 | MLE | MAGE |
| --- | --- | --- | --- | --- | --- | --- | --- | --- | --- |
| **JSD** | 0.8139 | 0.7337 | 0.8294 | 0.7441 | 0.7508 | 0.7074 | 0.7084 | 0.3188 | **0.3015** |
| **TVD** | 0.9588 | 0.8061 | 0.9982 | 0.8528 | 0.8442 | 0.8585 | 0.8599 | 0.2033 | **0.1560** |
| **HD** | 0.9776 | 0.8783 | 0.9892 | 0.8889 | 0.9002 | 0.8090 | 0.8110 | 0.3706 | **0.3606** |
| **CM** | 0.0436 | 0.2167 | 0.0018 | 0.3157 | 0.2235 | **0.9486** | 0.9338 | 0.8308 | 0.8930 |
| **Prec.** | 0.0714 | 0.0342 | 0.0262 | 0.0631 | 0.0471 | 0.0095 | 0.0093 | 0.2624 | **0.8772** |
| **Recall** | 0.0333 | 0.2833 | 0.3000 | 0.1167 | 0.1333 | 0.1667 | 0.1667 | 0.9667 | **0.8333** |
| **F1** | 0.0455 | 0.0610 | 0.0481 | 0.0819 | 0.0696 | 0.0180 | 0.0175 | 0.4128 | **0.8547** |

**Table 3.**
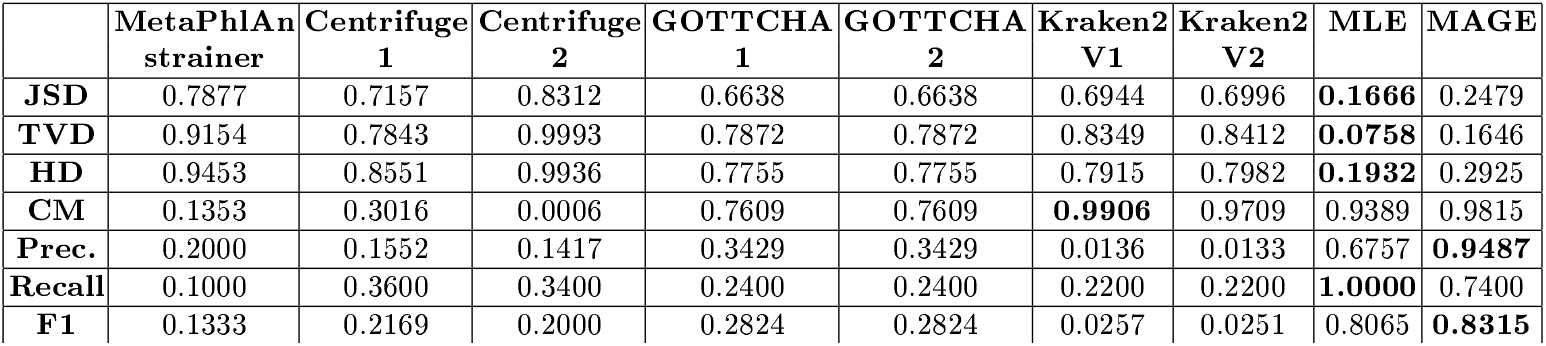
Strain level profiling performances on LC data

As seen in the tables, for the high complexity datasets (HC1 and HC2), MAGE achieves significantly higher F1 score and lower distances. We remark that higher recall values MLE comes at the cost of lower precision and significantly lower F1 due to the presence of several false positives. For the low complexity dataset (LC), MAGE achieves higher F1 score. For LC datasets, MLE achieved better distances, closely followed by MAGE.

### 3.5 Results on Strain Mutation Experiments

Table 4, Table 5 and Table 6 shows the comparative performances for HC1Mutated, HC2-Mutated and LC-Mutated datasets. MAGE exhibited superior performance on majority of the performance metrics.

**Table 4.** Strain level profiling performances on HC1 Mutated dataset

|  | MetaPhlAn<br>strainer | Centrifuge<br>1 | Centrifuge<br>2 | GOTTCHA<br>1 | GOTTCHA<br>2 | Kraken2<br>V1 | Kraken2<br>V2 | MLE | MAGE |
| --- | --- | --- | --- | --- | --- | --- | --- | --- | --- |
| **JSD** | 0.8170 | 0.7627 | 0.8308 | 0.6971 | 0.7527 | 0.6938 | 0.6958 | 0.5417 | **0.4626** |
| **TVD** | 0.9684 | 0.8805 | 0.9990 | 0.7714 | 0.8608 | 0.8158 | 0.8176 | 0.6088 | **0.4674** |
| **HD** | 0.9803 | 0.9084 | 0.9930 | 0.8328 | 0.9016 | 0.7908 | 0.7935 | 0.6276 | **0.5429** |
| **CM** | 0.0341 | 0.1475 | 0.0010 | 0.4205 | 0.1392 | **0.9686** | 0.9574 | 0.3912 | 0.5328 |
| **Prec.** | 0.1538 | 0.0437 | 0.0272 | 0.0795 | 0.0552 | 0.0101 | 0.0100 | 0.0667 | **0.4923** |
| **Recall** | 0.0588 | 0.2941 | 0.3235 | 0.2059 | 0.2353 | 0.2353 | 0.2353 | **0.9706** | 0.9412 |
| **F1** | 0.0851 | 0.0760 | 0.0502 | 0.1148 | 0.0894 | 0.0194 | 0.0191 | 0.1248 | **0.6465** |

**Table 5.** Strain level profiling performances on HC2 Mutated dataset

|  | MetaPhlAn<br>strainer | Centrifuge<br>1 | Centrifuge<br>2 | GOTTCHA<br>1 | GOTTCHA<br>2 | Kraken2<br>V1 | Kraken2<br>V2 | MLE | MAGE |
| --- | --- | --- | --- | --- | --- | --- | --- | --- | --- |
| **JSD** | 0.8029 | 0.7695 | 0.8295 | 0.7444 | 0.7532 | 0.7075 | 0.7085 | 0.5041 | **0.4754** |
| **TVD** | 0.9333 | 0.8771 | 0.9982 | 0.8523 | 0.8436 | 0.8598 | 0.8612 | 0.5191 | **0.3984** |
| **HD** | 0.9644 | 0.9214 | 0.9898 | 0.8896 | 0.9033 | 0.8086 | 0.8105 | 0.5869 | **0.5686** |
| **CM** | 0.0719 | 0.1328 | 0.0018 | 0.3099 | 0.2094 | **0.9567** | 0.9423 | 0.4954 | 0.6613 |
| **Prec.** | 0.0968 | 0.0286 | 0.0290 | 0.0593 | 0.0442 | 0.0083 | 0.0083 | 0.1473 | **0.3895** |
| **Recall** | 0.0500 | 0.2000 | 0.2667 | 0.1167 | 0.1333 | 0.1500 | 0.1500 | **0.9500** | 0.6167 |
| **F1** | 0.0659 | 0.0501 | 0.0524 | 0.0787 | 0.0664 | 0.0157 | 0.0157 | 0.2550 | **0.4774** |

**Table 6.** Strain level profiling performances on LC Mutated dataset

|  | MetaPhlAn<br>strainer | Centrifuge<br>1 | Centrifuge<br>2 | GOTTCHA<br>1 | GOTTCHA<br>2 | Kraken2<br>V1 | Kraken2<br>V2 | MLE | MAGE |
| --- | --- | --- | --- | --- | --- | --- | --- | --- | --- |
| **JSD** | 0.7764 | 0.7287 | 0.8315 | 0.6649 | 0.6649 | 0.6952 | 0.7003 | **0.2874** | 0.2990 |
| **TVD** | 0.8956 | 0.8054 | 0.9995 | 0.7872 | 0.7872 | 0.8359 | 0.8428 | 0.2148 | **0.1859** |
| **HD** | 0.9315 | 0.8710 | 0.9952 | 0.7775 | 0.7775 | 0.7924 | 0.7989 | 0.3366 | **0.3541** |
| **CM** | 0.1677 | 0.2656 | 0.0005 | 0.7482 | 0.7482 | **0.9917** | 0.9730 | 0.8022 | 0.8941 |
| **Prec.** | 0.1515 | 0.0057 | 0.0039 | 0.3077 | 0.3077 | 0.0132 | 0.0132 | 0.4673 | **0.8837** |
| **Recall** | 0.1000 | 0.3200 | 0.3000 | 0.2400 | 0.2400 | 0.2200 | 0.2200 | **1.0000** | 0.7600 |
| **F1** | 0.1205 | 0.0111 | 0.0077 | 0.2697 | 0.2697 | 0.0249 | 0.0250 | 0.6369 | **0.8172** |

### 3.6 MAGE Implementation

MAGE was implemented in C++ as multi-threaded. The R-Index library [11, 15] and FM-index library [24] were used for indexing level 2 sequence sub-collections. The total size of the MAGE index was 38GB. MAGE mapper takes around 1.9 hours using 64 threads for mapping a sample of 1 million reads.

## 4 Conclusion

Shotgun metagenomic sequencing has greatly empowered the exploration of metagenomic samples at a high resolution of strains. Current state-of-art methods predominantly focus on the problem of strain level identification and species level abundance quantification. Strain level abundance estimation is challenging and remains largely unaddressed. We develop MAGE (Microbial Abundance GaugE) which allows for deep taxonomic profiling of metagenome sample. It enables accurate estimation of strain level abundances which plays a crucial role in various downstream analyses. For accurate profiling, MAGE makes use of read mapping information and performs a novel local search-based profiling guided by a constrained optimization based on maximum likelihood estimation. Unlike the existing approaches which often rely on strain-specific markers and homology information for deep profiling, MAGE works solely with read mapping information, which is the set of target strains from the reference collection for each mapped read. As part of MAGE, we also provide an alignment-free and kmer-based read mapper that uses a compact and comprehensive index constructed using FM-index and R-index. We use a variety of evaluation metrics for validating abundances estimation quality. We have extensively benchmarked our results on low complexity and high complexity datasets, which demonstrates the ability of MAGE for accurate profiling. Our work shows that even in the absence of strain specific markers or marker guided read filtering, strain level profiling can be performed solely from the read mapping information. Unlike many of the existing approaches, no separate processing using various taxonomic markers is required for handling read mapping ambiguities. As a future work, it would be interesting to explore improved methods to guide the strain set refinement using more informative surrogates. We believe that our work would drive further research on novel approaches for abundance quantification at strain level resolution and various metrics for benchmarking.

## References

1. Alizon, S., de Roode, J.C., Michalakis, Y.: Multiple infections and the evolution of virulence. Ecology letters 16(4), 556–567 (2013)

2. Anyansi, C., Straub, T.J., Manson, A.L., Earl, A.M., Abeel, T.: Computational methods for strain-level microbial detection in colony and metagenome sequencing data. Frontiers in Microbiology 11, 1925 (2020)

3. Balmer, O., Tanner, M.: Prevalence and implications of multiple-strain infections. The Lancet infectious diseases 11(11), 868–878 (2011)

4. Beghini, F., McIver, L.J., Blanco-Míguez, A., Dubois, L., Asnicar, F., Maharjan, S., Mailyan, A., Manghi, P., Scholz, M., Thomas, A.M., et al.: Integrating taxonomic, functional, and strain-level profiling of diverse microbial communities with biobakery 3. Elife 10, e65088 (2021)

5. Bray, N.L., Pimentel, H., Melsted, P., Pachter, L.: Near-optimal probabilistic rna-seq quantification. Nature biotechnology 34(5), 525–527 (2016)

6. Centrifuge. https://ccb.jhu.edu/software/centrifuge/

7. Da Silva, K., Pons, N., Berland, M., Oñate, F.P., Almeida, M., Peterlongo, P.: Strainflair: strain-level profiling of metagenomic samples using variation graphs. PeerJ 9, e11884 (2021)

8. van Dijk, L.R., Walker, B.J., Straub, T.J., Worby, C.J., Grote, A., Schreiber, H.L., Anyansi, C., Pickering, A.J., Hultgren, S.J., Manson, A.L., et al.: Strainge: a toolkit to track and characterize low-abundance strains in complex microbial communities. Genome biology 23(1), 1–27 (2022)

9. Ferragina, P., Manzini, G.: Opportunistic data structures with applications. In: Proceedings 41st Annual Symposium on Foundations of Computer Science. pp. 390–398. IEEE (2000)

10. Freitas, T.A.K., Li, P.E., Scholz, M.B., Chain, P.S.: Accurate read-based metagenome characterization using a hierarchical suite of unique signatures. Nucleic acids research 43(10), e69–e69 (2015)

11. Gagie, T., Navarro, G., Prezza, N.: Optimal-time text indexing in bwt-runs bounded space. In: Proceedings of the Twenty-Ninth Annual ACM-SIAM Symposium on Discrete Algorithms. pp. 1459–1477. SIAM (2018)

12. Hamady, M., Knight, R.: Microbial community profiling for human microbiome projects: tools, techniques, and challenges. Genome research 19(7), 1141–1152 (2009)

13. Huang, W., Li, L., Myers, J.R., Marth, G.T.: Art: a next-generation sequencing read simulator. Bioinformatics28(4), 593–594 (2012)

14. Kim, D., Song, L., Breitwieser, F.P., Salzberg, S.L.: Centrifuge: rapid and sensitive classification of metagenomic sequences. Genome research 26(12), 1721–1729 (2016)

15. Kuhnle, A., Mun, T., Boucher, C., Gagie, T., Langmead, B., Manzini, G.: Efficient construction of a complete index for pan-genomics read alignment. Journal of Computational Biology 27(4), 500–513 (2020)

16. Li, H.: wgsim - simulating sequence reads from a reference genome. https://github.com/lh3/wgsim (2011)

17. Li, H., Handsaker, B., Wysoker, A., Fennell, T., Ruan, J., Homer, N., Marth, G., Abecasis, G., Durbin, R.: The sequence alignment/map format and samtools. Bioinformatics 25(16), 2078–2079 (2009)

18. Lu, J., Breitwieser, F.P., Thielen, P., Salzberg, S.L.: Bracken: estimating species abundance in metagenomics data. PeerJ Computer Science 3, e104 (2017)

19. McIntyre, A.B., Ounit, R., Afshinnekoo, E., Prill, R.J., Hénaff, E., Alexander, N., Minot, S.S., Danko, D., Foox, J., Ahsanuddin, S., et al.: Comprehensive bench-marking and ensemble approaches for metagenomic classifiers. Genome biology 18(1), 1–19 (2017)

20. McIver, L.J., Abu-Ali, G., Franzosa, E.A., Schwager, R., Morgan, X.C., Waldron, L., Segata, N., Huttenhower, C.: biobakery: ta metaâĂZomic analysis environment. Bioinformatics 34(7), 1235–1237 (2018)

21. MetaPhlAn2. https://github.com/biobakery/MetaPhlAn2

22. Nikulin, M.S., et al.: Hellinger distance. Encyclopedia of mathematics 78 (2001)

23. O’Leary, N.A., Wright, M.W., Brister, J.R., Ciufo, S., Haddad, D., McVeigh, R., Rajput, B., Robbertse, B., Smith-White, B., Ako-Adjei, D., et al.: Reference sequence (refseq) database at ncbi: current status, taxonomic expansion, and functional annotation. Nucleic acids research 44(D1), D733–D745 (2016)

24. Petri, M.: Fm-index-compressed full-text index. https://github.com/mpetri/FM-Index (2015)

25. Roberts, A., Pachter, L.: Streaming fragment assignment for real-time analysis of sequencing experiments. Nature methods 10(1), 71–73 (2013)

26. Scholz, M., Ward, D.V., Pasolli, E., Tolio, T., Zolfo, M., Asnicar, F., Truong, D.T., Tett, A., Morrow, A.L., Segata, N.: Strain-level microbial epidemiology and population genomics from shotgun metagenomics. Nature methods 13(5), 435–438 (2016)

27. Simon, H.Y., Siddle, K.J., Park, D.J., Sabeti, P.C.: Benchmarking metagenomics tools for taxonomic classification. Cell 178(4), 779–794 (2019)

28. Sims, G.E., Jun, S.R., Wu, G.A., Kim, S.H.: Alignment-free genome comparison with feature frequency profiles (ffp) and optimal resolutions. Proceedings of the National Academy of Sciences 106(8), 2677–2682 (2009)

29. Truong, D.T., Franzosa, E.A., Tickle, T.L., Scholz, M., Weingart, G., Pasolli, E., Tett, A., Huttenhower, C., Segata, N.: Metaphlan2 for enhanced metagenomic taxonomic profiling. Nature methods 12(10), 902–903 (2015)

30. Wood, D.E., Lu, J., Langmead, B.: Improved metagenomic analysis with kraken 2. Genome biology 20(1), 1–13 (2019)

31. Wood, D.E., Salzberg, S.L.: Kraken: ultrafast metagenomic sequence classification using exact alignments. Genome biology 15(3), 1–12 (2014)

